# Non-continuous neuromodulation in awake, unrestrained felines increases bladder capacity

**DOI:** 10.1101/2024.12.19.629379

**Authors:** Miguel Ortiz-Lopez, Zhonghua Ouyang, Anagha Kotkar, Maeve Willen, Richard Liu, Jordan Olszewski, Miriam Stevens, Eric Kennedy, Sarah Offutt, Katie Bittner, Lance Zirpel, Tim M. Bruns

## Abstract

**Objective:** Neuromodulation is a standard therapy for bladder symptoms such as overactive bladder. Previous studies have demonstrated that non-continuous stimulation (NCS) can increase bladder capacity and that bladder pressure can be estimated from dorsal root ganglia (DRG) neural activity in anesthetized animal models. Our goal is to determine if NCS elicits similar bladder capacity effects as continuous stimulation (CS) and if bladder pressure can be estimated from DRG signals in an awake, unrestrained animal model.

**Approach:** We performed aseptic, chronic implant surgeries with seven adult, male felines. Three animals were used to establish procedures, three for experimental testing, and one did not yield data. Bipolar stimulating electrodes were placed on the pudendal nerve and sacral nerve on the same side. Microelectrode arrays were inserted in two ipsilateral sacral DRG. Two single-lumen catheters were implanted in the bladder dome for recording bladder pressure and infusing saline. Fixed-sequence, repeated bladder fills were performed in four awake felines to evaluate the bladder capacity during no-stimulation (NS), NCS, and CS at either the pudendal or sacral nerve. NCS was performed based on increases in bladder pressure estimated from DRG recordings or when 50% of the average NS bladder capacity was reached.

**Main results:** We observed similar bladder capacity increases for NCS (122 ± 31% of NS control) as for CS (121 ± 33%) in the four animals. NCS paradigms reduced stimulation time by 46% on average. Median correlation coefficients of 0.46 and 0.64 (maximum 0.93) between the predicted and measured bladder pressure were obtained for awake trials with DRG bladder units in two animals.

**Significance:** This study demonstrated the feasibility of using NCS to increase bladder capacity in awake, unrestrained felines and for decoding bladder pressure from DRG recordings. Further studies are needed to optimize NCS timing for clinical translation.

## 1. Introduction

Overactive bladder (OAB) is a common dysfunction that affects women and men [1]. The prevalent symptom is urinary urgency sensation with or without incontinence, even when urine volume is low. The first line of treatment for OAB typically consists of behavioral changes such as pelvic floor muscle training and fluid intake modification [2]. Anticholinergic and β-adrenergic drugs are standard second-line treatments. However, many patients with OAB discontinue these treatments within one year due to poor efficacy or side effects [3]. Clinicians may offer chemodenervation via intradetrusor injection of OnabotulinumtoxinA (Botox), posterior tibial nerve stimulation (PTNS), or sacral neuromodulation (SNM) as third-line treatments [4]. In SNM, a four-contact stimulation lead is placed by a sacral nerve, typically S3. This spinal level includes primary innervation of the pelvic viscera and floor, including the bladder, urethral sphincter, and pelvic floor muscles, via the pelvic and pudendal nerves. Distal neuromodulation directly at the pudendal nerve, which is comprised of fibers from sacral nerves S2-S4, has also been studied for OAB and other bladder dysfunctions [5,6].

Some SNM patients have a relapse of symptoms or efficacy loss after a few years [7]. The default setting for the SNM device is continuous stimulation (CS) at 14 Hz [8]. In other applications, prolonged nerve exposure to CS can induce habituation [9]. Sacral nerve habituation may contribute to unsuccessful SNM outcomes, though it has not been proven. Clinical studies have demonstrated that cycling SNM (e.g. sixteen seconds on and eight seconds off) provides similar therapeutic efficacy to continuous SNM [8], suggesting that continuous SNM may not be necessary to elicit equivalent therapy efficacy. Pre-clinical and clinical studies have demonstrated that triggering pudendal or sacral stimulation on a specific part of the bladder filling cycle or during detrusor activity can increase bladder capacity [10–13]. This prior work suggests that stimulation that takes into consideration the bladder state could be a promising next step for the therapy and may prolong clinical outcomes by stimulating only when needed. However, further studies are required to compare the efficacy of CS with that of non-continuous stimulation (NCS), such as with bladder activity-related closed-loop stimulation (CLS).

Implementing a CLS system involves new challenges: identifying reliable biomarkers and developing algorithms to control stimulus timing. One potential biomarker are bladder sensory signals that can be observed with microelectrodes at sacral-level dorsal root ganglia (DRG). We previously demonstrated that DRG signals can be used to estimate bladder pressure [10,14] and as a biomarker for CLS in anesthetized settings [10,11]. The majority of prior neuromodulation studies targeting the lower urinary tract in animal models have been conducted in anesthetized subjects [15]. However, anesthesia is known to alter bladder and neural responses [16], limiting the clinical relevance of these findings. Moving beyond anesthetized studies is critical for translating neuromodulation therapies.

The primary objective of this study was to compare the effects of NCS and CS on the bladder in awake, healthy felines. The secondary objective was to estimate bladder pressure in real-time from sacral-level DRG recordings to trigger CLS in awake felines. We hypothesized that NCS would yield similar bladder capacity increases as CS but would significantly reduce the applied stimulation time, as has been observed under anesthesia [11].

## 2. Methods

Table 1 summarizes the goals for each animal. The first three animals (Animals 1-3) were used to establish the experimental procedures. Animals 1 and 2 were used to test the decoding algorithms in sedated test sessions. Animal 3 was used to establish awake experimental logistics and procedures for repeated test sessions. Finally, Animals 4 through 6 were used for repeated neuromodulation sessions to evaluate the bladder capacity during no-stimulation (NS), NCS, and CS at either the pudendal or sacral nerve. In this study, a trial is defined as a voiding cycle that begins with the start of infusion and ends with a voiding event.

**Table 1.** Summary of experiments performed in each animal (numbered 1-6).

| # | Goal(s) | DRG<br>Implant side<br>S1/S2 | DRG<br>bladder<br>units | Sacral<br>Neuromodulation | Pudendal<br>Neuromodulation | NCS<br>type | Litterbox<br>tracking |
| --- | --- | --- | --- | --- | --- | --- | --- |
|  |  |  |  | Sedated /<br>Awake testing | Sedated / Awake<br>testing |  |  |
| 1 | Bladder decoding<br>during sedation;<br>habituation<br>protocol | Left/Left | Yes | Yes / No | Yes / No | None | Yes |
| 2 | Bladder decoding<br>during sedation;<br>habituation<br>protocol | Left/Left | Yes | Yes/ Yes | No / No | Half<br>volume | No |
| 3 | Awake testing<br>logistics | Left/Right | Yes | Yes / Yes | No / Yes | Closed<br>loop | Yes |
| 4 | Repeated<br>neuromodulation | Left/Left | Yes | No / No | No / Yes | Closed<br>loop | Yes |
| 5 | Repeated<br>neuromodulation | Left/Left | No | No / Yes | No / Yes | Half<br>volume | Yes |
| 6 | Repeated<br>neuromodulation | Left/Left | No | No / Yes | No / Yes | Half<br>volume | Yes |

NCS – Non-continuous stimulation

### 2.1 Animals

All animal care and experimental procedures were approved by the University of Michigan Institutional Animal Care and Use Committee (IACUC, Protocol PRO00009298), following the National Institute of Health’s guidelines for the care and use of laboratory animals. Procedures were conducted in seven purpose-bred adult, neurologically-intact male felines (1.29 ± 0.19 years old, 4.80 ± 0.57 kg at surgery). Shortly after implant recovery, Animal 7 had a health issue related to the suprapubic catheter and subsequently was removed from the study. Animals were free-range housed in a 6.7 m by 6.4 m room with controlled temperature (19-21 °C) and relative humidity (35-60%), food and water available ad-lib, and a 12-hour light/dark cycle. Animals received enrichment via staff interaction and toys.

### 2.2 Surgical procedure

For each implant surgery, anesthesia was induced by an intramuscular administration of a mixture of Ketamine (6.6 mg/kg), Butorphanol (0.66 mg/kg), and Dexmedetomidine (0.011 mg/kg). Animals were intubated and transitioned to 1-4% isoflurane anesthesia during surgical procedures. Heart rate, O2 perfusion, temperature, and blood pressure were monitored, and an intravenous line was placed for saline infusion (10-20 ml/hr). After stabilizing the animals under anesthesia, a laparotomy was performed to insert two one-mm inner diameter Silastic catheters into the dome of the bladder. These catheters, intended for saline infusion and recording bladder pressure, were secured with a purse-string suture and tunneled to a midline incision in the lower back. Subsequently, a postero-lateral incision was used to access the left pudendal nerve to place a bipolar cuff electrode (2-mm diameter, 2-2.5 mm wire separation) for stimulation. The cuff electrode wires (32 AWG-AS 636, Cooner Wire, Chatsworth, CA) were also tunneled to the back incision. Following this, the skin and muscle were reflected in layers at the back incision to expose the L7-S3 vertebrae. The dorsal process and lamina of the S1 and S2 vertebrae were removed to expose the DRG. These spinal levels are equivalent to S2 and S3 in humans and are common SNM targets in feline studies [11,17]. Two wire electrodes (32 AWG-AS 636, Cooner Wire, Chatsworth, CA) in bipolar configuration were secured in place on the top of the right S1 spinal nerve just distal to the DRG for SNM with stay sutures secured to nearby muscle tissue. Microelectrode arrays (4×8 layout, IrOx 1 mm shanks, Blackrock Microsystems, Salt Lake City, UT) were inserted into the left S1 and left S2 DRG using a pneumatic inserter (Figure 1). For Animal 3, the second array was placed in the right S2 DRG. Finally, a four-post stainless-steel base plate was secured to the iliac crests below the skin, with the posts inserted through the skin. A 3D-printed chamber, or backpack, was attached to the baseplate posts and used to hold catheter ports and electrode connectors within secured locations [18,19]. After completing all surgical steps, standard post-op care and analgesia were provided while the animal recovered. Experimental testing was not performed for at least one week after surgery.

**Figure 1.**
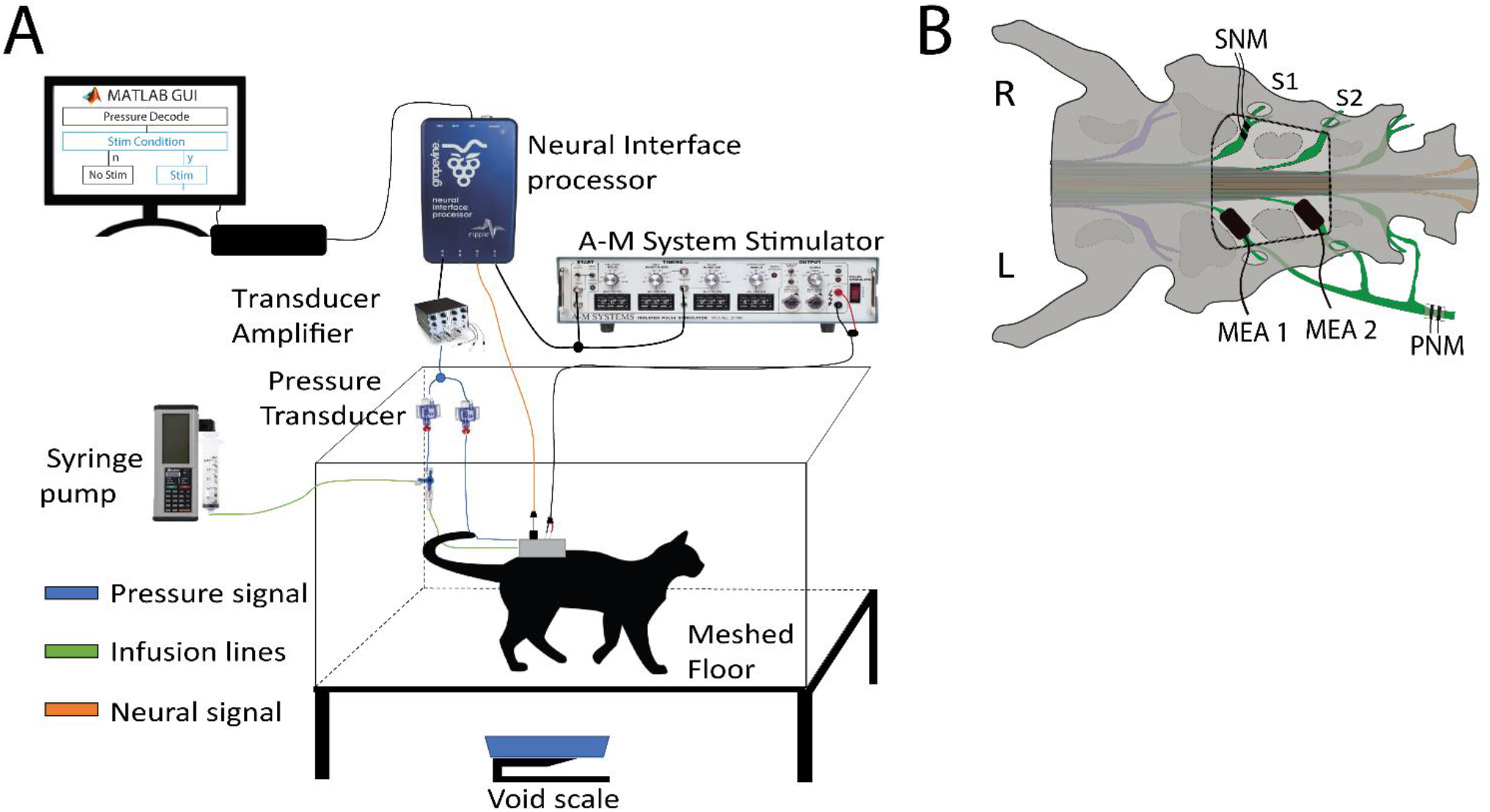
Experimental setup. A. Experimental test setup for awake, unrestrained sessions. The animal was placed in the test chamber and connected to the infusion pump, pressure transducers, data acquisition system, and the stimulator. B. Typical electrode placement following exposure of sacral DRG with a partial laminectomy between S1-S2 vertebrae. Two wire electrodes in bipolar configuration were placed on the top of the right S1 spinal nerve just distal to the DRG for SNM. Microelectrode arrays (MEA) were inserted into the left S1 and left S2 DRG. For Animal 3, MEA 2 was placed in the right S2 DRG. A bipolar cuff was placed on the left pudendal nerve via a separate surgical access.

### 2.3 Experiment setup and data collection

Bladder pressure was monitored with pressure transducers (DPT-100, Utah Medical Products, Midvale, UT) connected to each implanted catheter and an amplifier (TBM4M, World Precision Instruments, Sarasota, FL). The bladder pressure signal was recorded at 1 kHz, using a Grapevine Neural Interface Processer (NIP) (Ripple, Salt Lake City, UT). Neural data from the microelectrodes were sampled at 30 kHz and band-passed filtered (250 Hz to 7.5 kHz) with the NIP and Grapevine Trellis software. During each bladder infusion trial, 0.9% warm saline (36°C) was infused (two or five mL/min) into the bladder with a syringe pump (AS50 Infusion Pump, Baxter International, Deerfield, IL, or Model NE-1000, New Era Pump Systems, Inc., Farmingdale, NY). The stimulation electrodes were connected to a pulse stimulator (Model 2100 Isolated Pulse Stimulator, A-M Systems, Sequim, WA).

Unsorted DRG spikes were used for bladder pressure estimation for CLS trials. The spikes were detected using a dual-threshold crossing at 3–5.5 times the root-mean-square value of the signal on each microelectrode channel. The pressure signal from the non-infusion catheter and neural spike times were streamed to a previously-developed online Kalman filter decoder in a MATLAB Graphic User Interface (MathWorks, Natick, MA) [10,11]. We implemented a common-mode rejection approach to address artifacts due to electrical stimulation or motion. If spikes were observed in more than 70% of the channels simultaneously, they were considered artifacts and subsequently removed from the analysis. First, the decoder was trained on a NS bladder infusion, specific to the animal and the day of the experiment. The firing rates of threshold-crossings on each channel were calculated using a two-second window average and each was correlated to the two-second window average of the bladder pressure. The Kalman filter was trained using channels with a firing rate correlation coefficient (CC) to pressure greater than a threshold of 0.7. If no channel had a CC greater than 0.7, the threshold was decreased in 0.1 steps until a channel met the criteria. If no channel had a CC greater than 0.2, CLS was not performed. The Kalman filter generated a pressure estimate every two seconds.

### 2.4 Experimental sessions

#### 2.4.1 Sedated test sessions

Animals were sedated at regular intervals (once every one or two weeks). Dexmedetomidine IM (0.01-0.04 mg/kg) was used to sedate the animals prior to maintenance with a continuous intravenous infusion of Alfaxalone (24-250 μg/kg/min) [20] until the end of testing. The primary goal of a sedated test session was to evaluate stimulation electrode and DRG recording microelectrode functionality prior to awake testing. Motor thresholds (MT) for the sacral and/or pudendal nerve electrodes were obtained at one Hz and 210 μs pulse width, and strength-duration curves with other pulse widths were obtained in at least two sessions. Impedances at 1 kHz were measured for DRG microelectrodes. In Animals 1-4, bladder fills (two or five ml/min) were performed to assess DRG neural signals and train a bladder pressure decoding algorithm for CLS. The MT was defined as the lowest stimulation amplitude that induced anal sphincter twitching. Additional sedation sessions were performed for backpack inspection and cleaning as needed.

#### 2.4.2 Awake, unrestrained test sessions

For an awake, unrestrained test session, the bladder catheters and electrodes were either connected while awake or during an immediately preceding sedation session. Testing for awake sessions were completed in a clear-walled 40 x 70 x 50 cm plexiglass chamber (Figure 1). The chamber allowed felines to freely move around in a controlled space, with cables draped outside the chamber and a scale underneath to collect urine voided through the meshed floor. Awake, unrestrained test sessions were only completed if the animal did not show anxiety and continued to engage with treats, toys, or wet food after being connected.

#### 2.4.3 Preparatory test sessions

Across Animals, we developed procedures to habituate animals to the testing chamber and experimental paradigm. In Animal 2, we focused on decoder training, examined the transition from sedated to awake conditions, and evaluated parameters for triggering neuromodulation. In Animal 3, sacral and pudendal CS trials at or above MT were performed during test sessions which had concurrent bladder filling. Real-time estimation of bladder pressure from DRG recordings in different experimental conditions was also performed in Animal 3 to establish optimal stimulation and recording conditions for the following animals. Additionally, three awake sessions were performed in Animal 3. In Supplement Part 1 we discuss lessons learned from these experiments.

#### 2.4.4 Awake repeated neuromodulation sessions

In Animals 4-6, at least seven awake, unrestrained test sessions were performed for each animal. Each session consisted of at least three sequential bladder fills for each of three stimulation paradigms: no stimulation, NCS, and CS. The per-session objective was to first perform bladder fills with no stimulation to train the DRG-signal decoder prior to performing bladder fills with CLS and then CS. A fixed-order, non-randomized fill sequence was used to address potential stimulation carry-over effect on subsequent periods and a cumulative effect of stimulation. Due to failures of the DRG microelectrodes for two Animals, NCS was performed in two paradigms: using neural signals to perform CLS (Animals 3 and 4) or by initiating stimulation when the bladder reached an average of half capacity of the NS fills (Animals 5 and 6). This half-capacity threshold was rationalized by prior observations that non-voiding contractions become more frequent as bladder capacity exceeds 50% [11]. In CLS trials, nerve stimulation was applied for 15 seconds when the estimated pressure increased at least 5 cmH_2_O in a 4 second window. Stimulation was applied at five Hz and MT, with a 210 μs pulse width, settings previously shown to increase bladder capacity in anesthetized felines [21]. Feline behavior was observed for signs that nerve stimulation was uncomfortable (e.g. stop engaging with wet food or responding to sensation). The number of trials per session, stimulation location, and stimulation paradigm varied per animal due to electrode performance and/or animal behavior.

### 2.5 Monitoring for neuromodulation carry-over effect

A video camera system (TV-DVR104K, TRENDnet, Torrance, CA) was used to monitor the litterboxes of the feline housing room. For each animal, the number of visits to a litterbox per 24-hour period were counted prior to the surgery, during surgery recovery, and before and after each test session. Two animals were kept in a recovery cage after surgery, either to recover further from surgery or to isolate from other animals and did not have data collected during those periods.

### 2.6 Study completion

After completion of all testing for each animal, a final surgery was performed for a final assessment of implant functionality and to explant DRG microelectrode arrays, simulation electrodes, backpack, and bladder catheters. Animals were euthanized with a dose of intravenous sodium pentobarbital (36 mg/kg) while under deep isoflurane anaesthesia.

### 2.7 Data analysis

The primary data analysis focused on analyzing the effect of neuromodulation on bladder capacity during the repeated neuromodulation sessions of Animals 3-6. For each bladder fill, the bladder capacity was measured and normalized to the NS control group average in each session. The bladder capacity for each fill sequence was calculated as the greater of the amount infused and the voided urine. Residual fluid in the bladder after voiding, if the infused volume was greater than the voided volume, was not removed to avoid the potential for pulling a vacuum inside the bladder. Voiding efficiencies for each fill sequence were calculated by dividing the voided volume by the bladder capacity. The peak bladder pressure increase was calculated as the 1-second average of the maximum pressure during voiding minus the average pressure during the first two seconds of the trial. Bladder compliance for each fill was calculated as the ratio between the bladder capacity and the peak bladder pressure. A one-way ANOVA was used to compare the effects of the different stimulation paradigms on each data measure, followed by Tukey HSD post hoc test for intra-group comparisons. To control for the increased risk of Type I error due to multiple metrics being analysed, a Bonferroni correction was applied. A linear regression was applied to the average NS bladder capacities over time.

For bladder fills with neural decoding from DRG signals, the correlation coefficient (R) between the measured pressure and estimated pressure was calculated to evaluate the performance of the decoding algorithm. Neural recordings were sorted offline for Animals with functioning DRG microelectrodes during bladder fills. Bladder units were defined as having a firing rate correlation coefficient to the bladder pressure greater than 0.2. A Mann-Whitney Rank Sum Test was used to compare the bladder pressure and bladder unit firing rate correlation between sedated and awake sessions for these units.

Two offline decoding strategies were evaluated. In the first, each trial was decoded using the same channel and model as in online decoding, but with noiseless firing rates (derived from multiunit-sorted channels) that were rescaled to match the mean and range of the original training data, thereby minimizing performance loss from distributional shifts. In the second, representing an upper-bound scenario, a model was trained for each trial by pooling all previous trials in which bladder units were present on the same set of channels, and decoding performance was reported for these optimally trained models. For all tests, a significance level of 0.05 was used. When relevant, results are presented as mean ± standard deviation.

## 3. Results

We performed successful implant surgeries with six animals. Table 1 has a summary of the experiments.

### 3.1 Electrode functionality

#### 3.1.1 Stimulation electrodes

All animals were implanted with both sacral and pudendal nerve stimulation electrodes. During the experimental period, 3 out of 6 sacral electrodes and 5 out of 6 pudendal electrodes remained functional and were able to evoke visible anal twitching (Figure S1, Table S1). Several implanted electrodes failed within two to three weeks. These failures were typically due to an unintended strain placed on the wires during early animal handling, except for the sacral electrode in Animal 4, for which the wires withdrew under the skin shortly after the surgery.

Thresholds for evoking anal twitching increased slightly over time for most electrodes, with greater variability observed in sacral than pudendal electrodes (Figure S1A-B). The average in-surgery pudendal electrode threshold at 210 µs was 153.5 ± 92.6 μA, and the median value across all post-surgery sessions was 230 μA (100–660 μA range). The average in-surgery sacral electrode threshold was 137.3 ± 98.5 μA and the median value across all post-surgery sessions was 127 μA (70–4102 μA range). Chronaxies for strength duration curves obtained from pudendal nerve and sacral nerve electrodes varied over time (Figure S1C-D).

#### 3.1.2 DRG recording microelectrodes

We observed DRG signals from four out of five Animals that had awake sessions. In Animal 6, the microelectrode cables were accidentally cut during the first test session due to poor surgical routing. Figure S2 summarizes microelectrode functionality and impedances after implant for other Animals.

### 3.2 DRG bladder units

Bladder units were detected during awake, unrestrained activities in three animals in offline analysis (Animals 2-4), with other units observed in four total Animals (Table S2). Figure S3 shows three representative bladder units during a NS bladder fill in Animal 3. Animals 3 and 4 had average bladder unit correlations of 0.41 ± 0.14 (range 0.20-0.74) and 0.49 ± 0.21 (0.21-0.92), respectively, between bladder unit firing rates and bladder pressure. A wide variability in sorted unit correlation to pressure and duration of tracking was observed across Animals 3 and 4 (Figure S4). Each Animal had one unit that was identifiable for the duration of testing, with considerable variability in its correlation over time. There were no differences in the firing characteristics of bladder units during NS and CS trials in Animal 4 (Figure S5), while Animals 2 and 3 did not have sufficient trials for a comparison.

### 3.3 Awake bladder pressure decoding

We utilized DRG recordings to estimate bladder pressure during awake bladder fills in Animals 3 and 4. Figure 2A & B show a representative NS trial used to train a bladder pressure model and an example pressure decoding trial in Animal 4. In Animal 3, there was high variability in the number of specific channels identified for decoding across sessions and trials (Figure 3A). For 38 total real-time decoding trials across 11 sessions, the average correlation coefficient was 0.45 ± 0.43 (median: 0.51, range: -0.63 – 0.90). Thirty of these trials used a single channel for the decode, and a total of 26 different channels were used at least once. Decoding of the bladder pressure during five ml/min bladder fills (correlation coefficient = 0.33 ± 0.45, n = 23) was lower (p = 0.03) than for two ml/min fills (0.63 ± 0.33, n = 15) in Animal 3.

**Figure 2.**
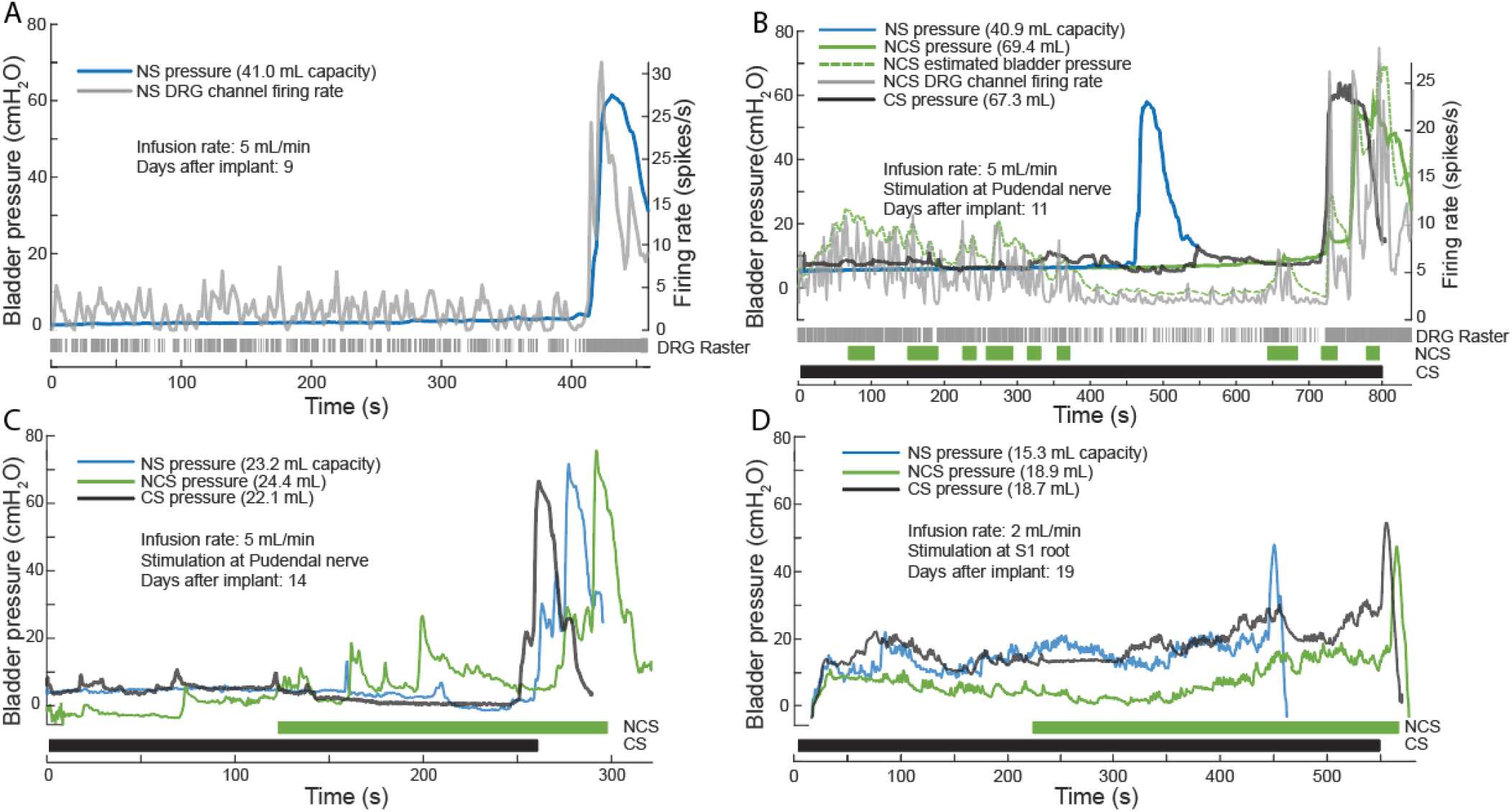
Representative trials across three Animals with repeated testing. A. NS bladder fill in Animal 4 used to train a bladder pressure model from one DRG channel. B. Representative stimulation trials in Animal 4, with the CLS trial using model of part A. C. Representative trials from Animal 5. D. Representative trials from Animal 6. (NS – No Stimulation, NCS – Non continuous Stimulation, CS – Continuous Stimulation)

**Figure 3.**
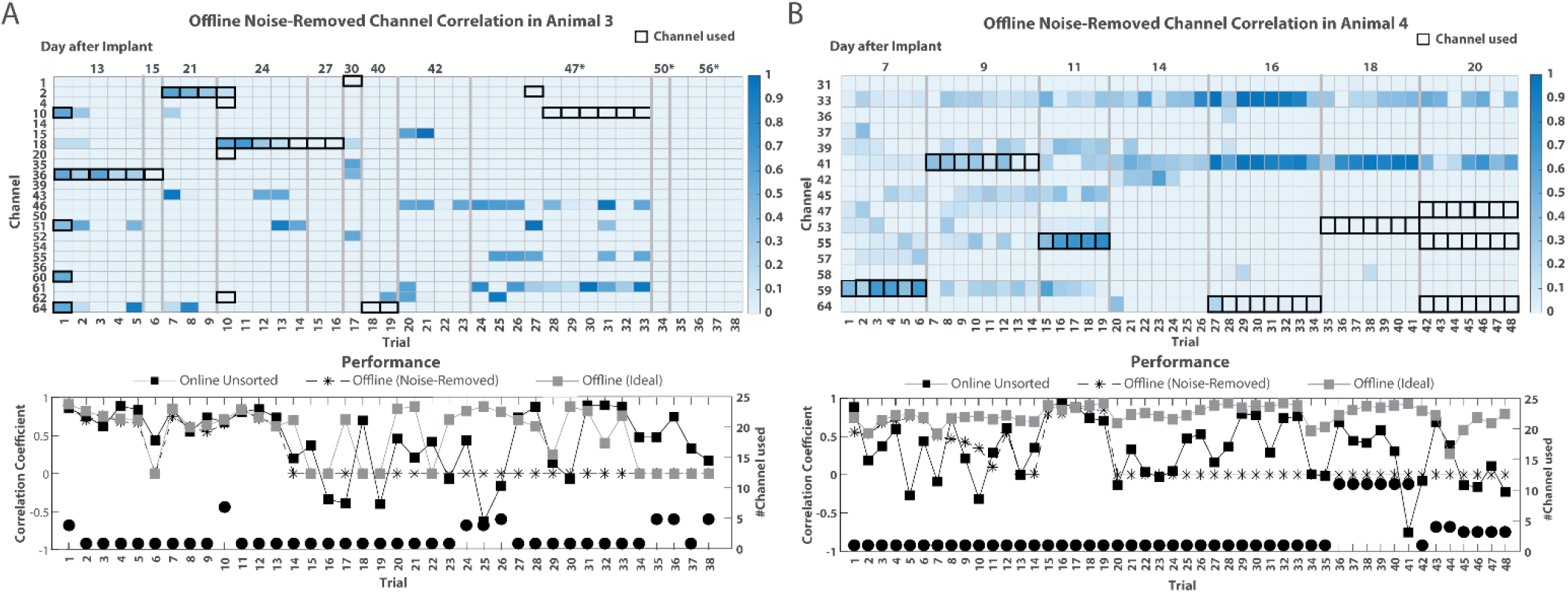
Offline decoding performance for unsorted channels with noise removed across awake trials in A. Animal 3 and B. Animal 4. Top panels: Correlations between firing rate and bladder pressure across channels and trials. Bolded squares represent the channels used during offline decoding. Not all channels are shown. Asterisks (*) denote days when the algorithm was trained during awake testing. Bottom panels: Bladder pressure decoding performance per trial. Black squares indicate online decoding performance. Asterisks indicate offline decoding results that match online decoding. Grey squares indicate ideal decoding performance. Black circles indicate the number of channels used for online decoding in each trial.

In Animal 4, the decoding performance had an overall lower performance (Figure 3B). For 48 total real-time decoding trials (28 CLS trials, 18 NS trials, and two CS trials when real-time decoding was accidentally left on) across seven sessions, the average correlation coefficient was 0.32 ± 0.39 (median: 0.39, range: -0.75 – 0.93). Thirty-five of these trials used a single channel for the decode, and a total of 11 channels were used at least once. Offline analysis showed that the bladder units identified during sedated trials were not always present during awake bladder fills. These inconsistencies resulted in some models being trained on channels without bladder units present during awake bladder fills, leading to decoding driven by noise or neural signals unrelated to bladder pressure (n = 22 trials in Animal 3; n = 29 trials in Animal 4). For awake trials in which real-time decoding was performed on channels with at least one bladder unit present, Animal 4 had a correlation coefficient of 0.44 ± 0.40 (median: 0.44, range: -0.31 – 0.93, n = 19), while Animal 3 had a higher and more consistent performance with a correlation coefficient of 0.64 ± 0.27 (median: 0.74, range: 0.14 – 0.90, n = 16). Offline decoding of sorted bladder units resulted in a mean correlation coefficient of 0.52 ± 0.32 (median: 0.56, range: 0.0–0.90, n = 16) and 0.55 ± 0.26 (median: 0.56, range: 0.09–0.88, n = 20) for Animals 3 and 4, respectively.

### 3.4 Awake repeated neuromodulation sessions

We performed DRG-based CLS in Animals 3 and 4. FigureB shows a set of CLS, CS, and NS trials in an Animal 4 session. In Animal 3, pudendal nerve CLS significantly increased bladder capacity (118 ± 7% of control capacity, Table 2, Figure 4A, p < 0.0125). In this animal CLS trials had an average duty cycle of 43 ± 17% and 83% of the stimulation was applied after half of the NS bladder capacity. In Animal 4, CS (144 ± 24% of control capacity) and pudendal nerve CLS (131 ± 21%) significantly increased bladder capacity (Table 2, Figure 4A, both p < 0.0125). The average duty cycle was 50 ± 21% in CLS trials and 72% of stimulation was applied after half of the NS bladder capacity in Animal 4.

**Table 2.**
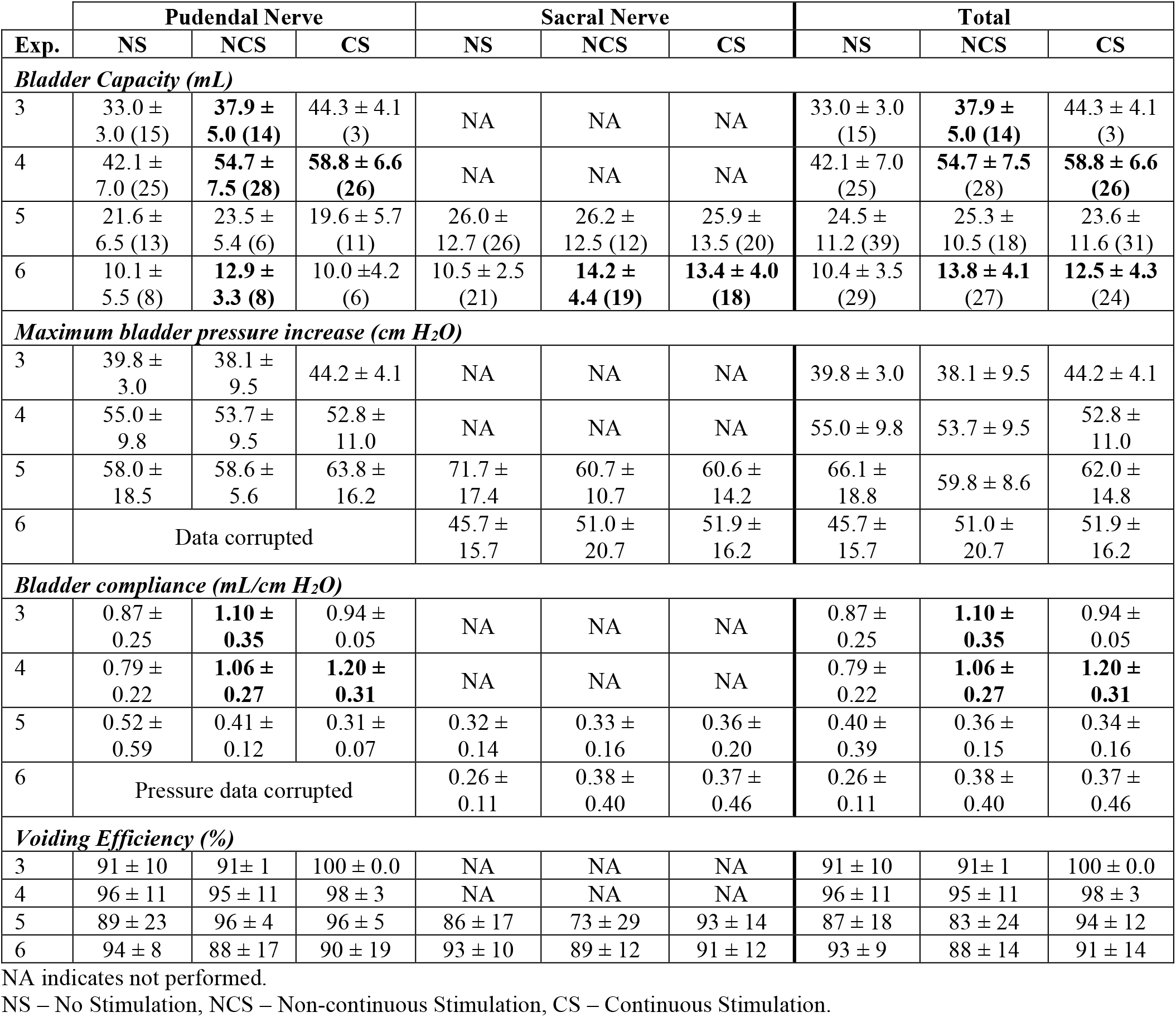
Summary of Repeated Neuromodulation Sessions across Animals 4-6. Sample sizes per Animal, stimulation location, and trial type are given as (N). Bold values are significantly different (p < 0.0125 from NS. Total is the average of all trials performed in an Animal – pudendal and sacral trials.

|  | Pudendal Nerve |  |  | Sacral Nerve |  |  | Total |  |  |
| --- | --- | --- | --- | --- | --- | --- | --- | --- | --- |
| Exp. | NS | NCS | CS | NS | NCS | CS | NS | NCS | CS |
| Bladder Capacity (mL) |  |  |  |  |  |  |  |  |  |
| 3 | 33.0 ± 3.0 (15) | 37.9 ± 5.0 (14) | 44.3 ± 4.1 (3) | NA | NA | NA | 33.0 ± 3.0 (15) | 37.9 ± 5.0 (14) | 44.3 ± 4.1 (3) |
| 4 | 42.1 ± 7.0 (25) | 54.7 ± 7.5 (28) | 58.8 ± 6.6 (26) | NA | NA | NA | 42.1 ± 7.0 (25) | 54.7 ± 7.5 (28) | 58.8 ± 6.6 (26) |
| 5 | 21.6 ± 6.5 (13) | 23.5 ± 5.4 (6) | 19.6 ± 5.7 (11) | 26.0 ± 12.7 (26) | 26.2 ± 12.5 (12) | 25.9 ± 13.5 (20) | 24.5 ± 11.2 (39) | 25.3 ± 10.5 (18) | 23.6 ± 11.6 (31) |
| 6 | 10.1 ± 5.5 (8) | 12.9 ± 3.3 (8) | 10.0 ± 4.2 (6) | 10.5 ± 2.5 (21) | 14.2 ± 4.4 (19) | 13.4 ± 4.0 (18) | 10.4 ± 3.5 (29) | 13.8 ± 4.1 (27) | 12.5 ± 4.3 (24) |
| Maximum bladder pressure increase (cm H<sub>2</sub>O) |  |  |  |  |  |  |  |  |  |
| 3 | 39.8 ± 3.0 | 38.1 ± 9.5 | 44.2 ± 4.1 | NA | NA | NA | 39.8 ± 3.0 | 38.1 ± 9.5 | 44.2 ± 4.1 |
| 4 | 55.0 ± 9.8 | 53.7 ± 9.5 | 52.8 ± 11.0 | NA | NA | NA | 55.0 ± 9.8 | 53.7 ± 9.5 | 52.8 ± 11.0 |
| 5 | 58.0 ± 18.5 | 58.6 ± 5.6 | 63.8 ± 16.2 | 71.7 ± 17.4 | 60.7 ± 10.7 | 60.6 ± 14.2 | 66.1 ± 18.8 | 59.8 ± 8.6 | 62.0 ± 14.8 |
| 6 | Data corrupted |  |  | 45.7 ± 15.7 | 51.0 ± 20.7 | 51.9 ± 16.2 | 45.7 ± 15.7 | 51.0 ± 20.7 | 51.9 ± 16.2 |
| Bladder compliance (mL/cm H<sub>2</sub>O) |  |  |  |  |  |  |  |  |  |
| 3 | 0.87 ± 0.25 | 1.10 ± 0.35 | 0.94 ± 0.05 | NA | NA | NA | 0.87 ± 0.25 | 1.10 ± 0.35 | 0.94 ± 0.05 |
| 4 | 0.79 ± 0.22 | 1.06 ± 0.27 | 1.20 ± 0.31 | NA | NA | NA | 0.79 ± 0.22 | 1.06 ± 0.27 | 1.20 ± 0.31 |
| 5 | 0.52 ± 0.59 | 0.41 ± 0.12 | 0.31 ± 0.07 | 0.32 ± 0.14 | 0.33 ± 0.16 | 0.36 ± 0.20 | 0.40 ± 0.39 | 0.36 ± 0.15 | 0.34 ± 0.16 |
| 6 | Pressure data corrupted |  |  | 0.26 ± 0.11 | 0.38 ± 0.40 | 0.37 ± 0.46 | 0.26 ± 0.11 | 0.38 ± 0.40 | 0.37 ± 0.46 |
| Voiding Efficiency (%) |  |  |  |  |  |  |  |  |  |
| 3 | 91 ± 10 | 91± 1 | 100 ± 0.0 | NA | NA | NA | 91 ± 10 | 91± 1 | 100 ± 0.0 |
| 4 | 96 ± 11 | 95 ± 11 | 98 ± 3 | NA | NA | NA | 96 ± 11 | 95 ± 11 | 98 ± 3 |
| 5 | 89 ± 23 | 96 ± 4 | 96 ± 5 | 86 ± 17 | 73 ± 29 | 93 ± 14 | 87 ± 18 | 83 ± 24 | 94 ± 12 |
| 6 | 94 ± 8 | 88 ± 17 | 90 ± 19 | 93 ± 10 | 89 ± 12 | 91 ± 12 | 93 ± 9 | 88 ± 14 | 91 ± 14 |

NA indicates not performed.

**Figure 4.**
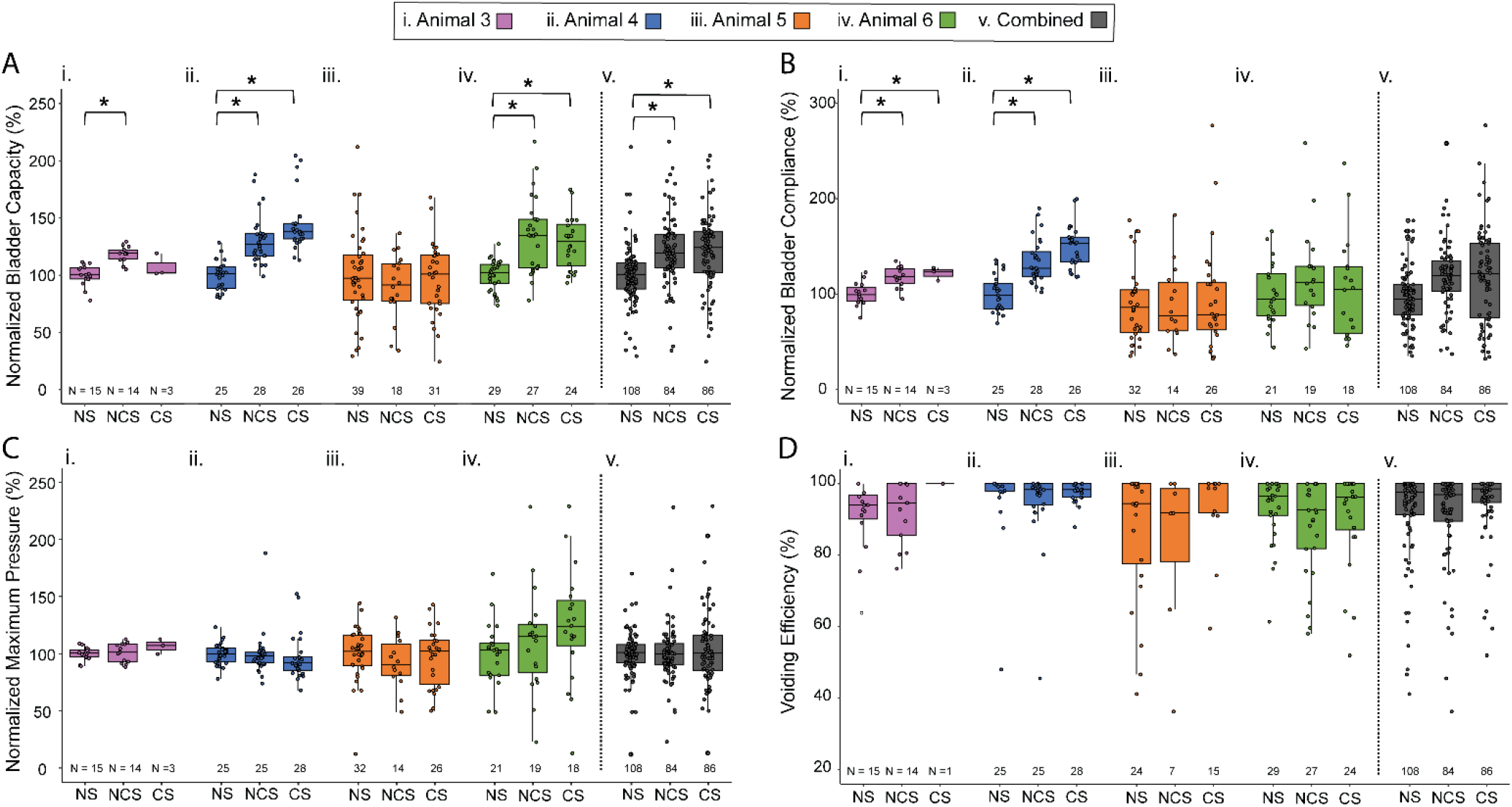
Normalized data measures across repeated neuromodulation sessions. Animals 3 and 4 received NCS using DRG-driven closed-loop control, while NCS in Animals 5 and 6 was initiated at a bladder volume threshold. A. Normalized bladder capacity. B. Normalized bladder compliance. C. Normalized maximum bladder pressure change. D. Voiding efficiency. Animals i. 3. ii. 4. iii. 5. iv. 6. v. Animals 3-6 combined. Boxes represent the interquartile range (IQR), with the line in the middle of each box representing the median. Whiskers extend to the smallest and largest data points that are within 1.5 times the IQR from the first and third quartiles, respectively. (* p < 0.0125 after Bonferroni correction, NS – No Stimulation, NCS – Non continuous Stimulation, CS – Continuous Stimulation).

In Animals 5 and 6 NCS was performed using a volume threshold. FigureC & D show representative trials for Animal 5 and 6, respectively. In Animal 5, stimulation was not observed to affect capacity, as NCS (90 ± 28%) and CS (96 ± 31%) across both nerve locations did not yield different capacities than NS trials. In Animal 6, pudendal nerve NCS (148 ± 46 %) and CS (128 ± 26 %) significantly increased bladder capacity. Similar bladder increases were achieved using sacral nerve NCS (129 ± 24 %) and CS (128 ± 22 %, both p < 0.0125). The average duty cycle for NCS trials was 46 ± 20% and 63 ± 17% for Animals 5 and 6, respectively. No significant differences in stimulation efficacy were observed between the sacral and pudendal nerve locations (Table 2).

Post hoc comparisons with a Tukey HSD test, followed by Bonferroni correction for multiple comparations across different metrics, identified statistically significant differences in bladder capacity during Animals 3, 4, and 6 and across all four Animals pooled together (Table 2, Figure 4, p < 0.0125). There were also significant differences in bladder compliance in Animals 3 and 4, but not across all Animals (Table 2, Figure 4, p < 0.0125). In contrast, no significant differences in peak bladder pressure or voiding efficiency were observed within animals or when data were pooled across animals.3.5 Stimulation effect on bladder function outside of sessions

We did not observe a carry-over effect of stimulation sessions on bladder capacity, based on tracking litterbox usage (Figure S6A, Table S3). In general, the animals had a consistently low count of litterbox visits prior to surgery, which increased in the recovery period after surgery. During weeks with experimental sessions, litterbox visits were significantly higher across animals than before surgery (p < 0.001), however there were no differences between pre-session and post-session visit counts within or across animals. The increase in litterbox visit counts varied across animals, from a 1.7x increase (Animal 4) up to an 8.6x increase (Animal 1) and there was not a trend of pre- or post-session measures being higher across animals (Table S3). Across Animals 4 through 6, there was not a cumulative increase in bladder capacity across the test sessions. While Animal 4 had a small, non-significant capacity increase over time, Animals 5 and 6 each had a linear decrease (p < 0.01) in bladder capacity across sessions (Figure S6B).

## 4. Discussion

In this study, we showed that, across the four animal tested, NCS (volume-threshold NCS or decoding-based CLS) can yield similar bladder capacity as CS in an awake, unrestrained feline model. Furthermore, we demonstrated the feasibility of decoding bladder pressure from DRG recordings in two animals. To our knowledge, this is the first study to perform real-time CLS using neural recordings in an awake, unrestrained medium-sized animal model towards restoring bladder function. This animal model provides a closer representation of the clinical setting than the standard use of anesthetized animals. Clinical and pre-clinical studies have shown that peripheral nerve NCS during non-voiding contractions or latter parts of a bladder fill can significantly increase bladder capacity [11,22–25]. In our study, neuromodulation was effective in three of four animals (Figure 4, Table 2). In the four animals, pudendal nerve NCS (volume-threshold NCS or decoding-based CLS) elicited a similar effect on bladder capacity as CS (Table 2), which aligns with a prior study with anesthetized animals [24].

Stimulation was also performed on the sacral nerve in two of these animals. In contrast to some clinical studies [5,26], we did not observe differences between the stimulus location and efficacy of stimulation (Table 2). The duration of stimulation in NCS trials for responding test animals averaged 46% less than for CS, with capacity increased by 30% compared to control (Figure 4,Table 2). These results are similar to a previous study with subjects with a spinal cord injury that detected bladder contractions with a urethral catheter to control dorsal genital nerve stimulation. In that study, CS and NCS increased bladder capacity by 36 ± 24% and 51 ± 37% compared to control, respectively, with conditional stimulation decreasing the stimulation time by 27 ± 17% [22]. Pre-clinical studies in anesthetized animals have shown that stimulation timing plays an essential role in therapy efficiency; optimal stimulation timing may be during non-voiding bladder contractions [23] or in the second half of bladder fills [25]. There is a correlation between these timings, as non-voiding bladder contractions became more frequent above half-bladder capacity [11]. Here, we obtained bladder capacity increases for stimulation during estimated bladder pressure increases (e.g., Figure 2B) and for fixed periods approaching a full bladder (e.g., Figure 2D). Further studies investigating optimal stimulation timings and the relevance of stimulation paradigms across sedated and awake animal models are needed.

Across awake sessions, stimulation was performed at MT and there was no sign of pain or anxiety. However, when stimulation was turned on some animals groomed the pelvic area, indicative of perception. At MT, we did not observe a decrease in voiding efficiency during stimulation trials (Table 2, Figure 4). The peak pressure during the voiding phase was not significantly different within or across animals though there were differences in compliance across responding animals. The significant increase in compliance with stimulation for Animals 3 and 4 (Figure 4B) aligns with a clinical neuromodulation study that observed detrusor inhibition without changes in urethral resistance (no differences in peak pressure) or detrusor contractility during the voiding phase, which resulted in higher bladder compliance [27]. However, in Animal 6, there were no significance changes in bladder compliance (Figure 4B).

In contrast to studies with anesthetized animals [10,11], we were unable to consistently decode bladder pressure from unsorted DRG spikes in awake animals (Figure 3). The primary factor was noise during awake recordings. The noise was typically caused by movement of the backpack and cables during position changes, leading to induced noise on the recordings. Spike detection using dual-threshold crossings with common-mode rejection was not sufficient to consistently reject noise. Another factor that affected our decoding performance was that units were not consistently present between sedated and awake testing, as we determined from offline spike sorting analysis. This variation led to channels selected for decoding for awake-testing bladder trials that often did not contain bladder-related activity, contributing to the high variability and low correlation coefficient in bladder pressure decoding (Figure 3). We defined a “bladder unit” as a channel whose firing rate correlated with bladder pressure with a correlation coefficient greater than 0.2, consistent with criteria used in anesthetized preparations [10]. This threshold has not been systematically evaluated in awake, unrestrained recordings, which is a limitation of the present study.

We hypothesize that microelectrode micromotion had a role in the variability of the correlation for tracked bladder units (Figure S4). Also, our use of sedated bladder fills to train the decoding algorithm, while assuming a preserved linear relationship between firing rate and bladder pressure across sedated and awake states, may have overfit the model. During the preparatory animals we evaluated using awake neural recordings for decoding, however they were inconsistent, foreshadowing our challenges in the test animals. Ultimately, model training during non-sedated bladder fills will be necessary to provide the most realistic scenario for translation of this approach. Also, reducing or eliminating recording noise during awake sessions will be essential for maintaining the spike detection algorithm as simple as possible. A sophisticated spike detection algorithm that considers waveform properties may be needed to reject noise [28,29]. An adaptive decoder [30] that uses data from multiple bladder fills and incorporates bladder neuron response characteristics, potentially capturing more complex relationships beyond linear correlations may also increase the accuracy of the decoder by accommodating the variability of the neural recordings [31,32].

We used microelectrodes that are not specifically designed for DRG, as in prior studies [10,11]. DRG units were observed for most animals (Table S1) and could be tracked over time across sedated and awake sessions (Figure S4), however functionality slightly decreased over time (Figure S2B) and we did not observe bladder units in all animals (Table S1). Mounting these rigid microelectrode arrays to small, curved DRG can be challenging, leading to arrays shifting over time [18] which may have contributed to the instability of sorted units (Figure 3). Flexible electrode arrays that can conform to the curved DRG surface [33] may have better success for identifying and tracking units over time. Alternatively, evoked responses that are recorded on the electrode lead may allow for a single electrode for CLS, as has been done for spinal cord stimulation [34].

In DRG-based CLS trials, nerve stimulation was applied for 15 seconds when the estimated bladder pressure increased by at least five cm H2O within a four-second window. This stimulation protocol assumes that non-voiding contractions, characteristic preclinical models of OAB, are present. In preclinical models, researchers typically induce hypersensitivity or bladder inflammation to study OAB and its treatments [35]. OAB symptoms are elicited through the intravesical infusion of external substances like acetic acid [15]. These methods reduce the bladder capacity and sometimes increase the frequency of non-voiding contractions during bladder fills. In this study, we used healthy felines and did not directly seek to create OAB conditions. However, most animals had large increases in their rate of litterbox usage after the implant surgery (Table S3, Figure S6A), suggesting an increase in urgency-like sensations. Additionally, two animals had a significantly decreasing trend over time in their bladder capacity (Figure S6B), which may have been caused by mechanical irritation of the implanted bladder catheters. Contrary to findings from sedated preclinical OAB models [10] and sedated bladder fills in this study, the pressure traces during awake session bladder fills did not show non-voiding contractions (Figure 2). This discrepancy raises questions regarding the relevance of the stimulation protocol in awake animals and whether the sedated model is a valid model for OAB. Future studies should examine the presence of non-voiding contractions during awake bladder fills to ascertain whether the stimulation algorithm used here remains applicable. Thus, in this study, the effects of stimulation were not a result of inhibiting non-voiding contractions; rather, the stimulation may have mitigated the effects of irritation during sessions.

In this study, stimulation carry-over effects were not observed. Bladder capacity increases were only observed during stimulation within sessions. This contrasts with clinical studies, which have shown maintained improvements in bladder symptoms with intermittent neuromodulation (eight hours on and sixteen hours off) [36] or sustained effects for days after cessation of neuromodulation [37]. Those carry-over effects were present after longer post-implant stimulation periods, suggesting that neuromodulation may need to be applied for a longer induction period before persistent effects are observed. In contrast with a prior study which suggested that bladder units can desensitize during CS in sedated bladder fills [11], our results did not have a consistent sensitivity trend during awake bladder fills with and without stimulation (Figure S5). Unfortunately, we did not perform sedated bladder fills with CS for a direct comparison to the prior study. A study with an implanted stimulator that can apply CS or NCS in an animal’s housing space, such as previously developed with sheep [38], may provide further insights into long-term efficacy and carry-over effects.

Neuromodulation to control bladder function in pre-clinical models is generally performed under anaesthesia [15]. Awake testing with rodent models is growing in prevalence, facilitated by the relative ease of tunnelling implant catheters and wires to connectors mounted on the skull, wireless stimulators and metabolic cages suited to rodent space requirements, and immobilization techniques that are better tolerated by small species [39,40]. Awake, unrestrained testing with medium-sized animals like felines brings new challenges (discussed in part in Supplement Part 1), including stress induced by handling, connecting implant monitoring system to highly mobile animals, and performing bladder fills in a controllable yet comfortable space. Several limitations of this study include a low sample size, a lack of stimulation type randomization within stimulation sessions and variations in animal behavior during trials (e.g., eating, walking or playing with toys). In addition, the use of two distinct non-continuous stimulation approaches represents a methodological consideration that may contribute to variability in the pooled analysis. Longer-term studies with untethered implants that utilize the animals’ natural bladder filling may allow for improved study design.

## 5. Conclusion

This study demonstrated that NCS can yield observable, significant bladder capacity increases without affecting peak pressures or voiding efficiency in a pre-clinical awake animal model. There is a clear promise for clinical use of NCS or even CLS; advances in recording electrode interfaces or approaches are needed. DRG may be a promising location for monitoring bladder pressure-related signals, as indicated by the large number of bladder units identified across Animals 2-4. We expect an electrode array specifically designed for DRG will have better long-term performance and help drive this therapeutic approach towards clinical utility. Future studies investigating NCS at different intervals during bladder fills are needed to maximize the therapy outcome while minimizing the energy required. Future studies with improved DRG interfaces that monitor bladder activity using complex decoders under awake, urestrained session and without test chambers are needed.

## Acknowledgments

The authors thank Elizabeth Bottorff, Po-Ju Chen, Amador Lagunas, Lauren Madden, Alexa Rybicki, and Joseph Wendt for assistance with data collection or analysis and the University of Michigan Unit for Laboratory Animal Medicine for assistance with animal care. A research grant from Medtronic supported this study. Katie Bittner, Sarah Offutt and Lance Zirpel were employees of Medtronic during this study. The opinions expressed in this article are the authors’ own and do not reflect the view of Medtronic.

