## Supplemental for "Non-continuous neuromodulation in awake, unrestrained felines increases bladder capacity"

#### Part 1- Experimental lessons learned

The following section describes challenges we faced during studies with initial Animals and improvements we made to experimental steps to address many of the challenges.

For sedation anesthesia, we determined that intramuscular dexmedetomidine (dex) could be used as a knockdown agent for subsequent intravascular alfaxalone, an anesthetic agent that can preserve bladder contractions better than other conventional agents [1]. We attempted oral dexmedetomidine (dex) in Animal 1 to reduce handling stress from intramuscular injections however we stopped using it due to its long time to take effect and the stress on the animal in this waiting period. Animals 2 and 3 tolerated intramuscular dex injections, assisted by close interactions between these felines and research staff. This trend continued across following Animals. Alfaxalone was given through a fresh intravascular catheter during each sedated session. This process was not an issue except that tissue scarring made catheterization a little more difficult over time. In this study all sedated bladder fills were performed under alfaxalone, although we performed some dex-only sedated sessions to perform tasks such as taking impedances, collecting strength-duration curves, and connecting to the implants.

In the first three Animals, we attempted to fine-tune the alfaxalone dosage to obtain strong bladder contractions for algorithm training; however, the results were not consistent. Empirically, we found whether bladder contractions occurred or not was more animal-dependent than dosage-dependent. We did not find strong evidence that a lighter alfaxalone dosage is more likely to allow strong bladder contractions while the animal was fully sedated, although we did notice strong bladder contractions after the animals were removed from alfaxalone and were half-awake. We also noticed a large residual volume while the animals were under alfaxalone sedation. Our conclusion from these alfaxalone tuning attempts was that alfaxalone sometimes preserved bladder contractions, which could lead to successful algorithm training, but dosage seemed to affect the plane of anesthesia more than bladder function. We do not recommend using a very low alfaxalone dosage in the hope of preserving awake-like bladder contractions, as it may not be a deep enough plane of anesthesia, leading to disruptions in the overall experimental flow.

We made progress with animal handling across the first three Animals that benefited testing across all animals. From the first day of arrival, animal handling was improved by regular interaction with staff, positive reinforcement, and associating negative events (mostly injections, pills, and awake experiments) with rewards (treats, wet food, and toys). We began by relying on sedation to connect to implants for awake testing and gradually increased the number of awake-only sessions from 0 (Animal 1) to 3 (Animal 2) and 4 (Animal 3) as we improved our methods for interacting with the animals. However, sedation before awake testing remained a reliable and low-risk option to connect to the implants. Each animal was conditioned to the testing room prior to surgery (one hour per session, at least four sessions), during which they were allowed to explore and were handled by study team members. After an implant surgery we continued conditioning the animals to the study team and test room with one week of daily one-hour conditioning sessions. We provided additional conditioning to the testing chamber depending on the need per animal. Animal 1 had awake testing in an open space rather than the chamber and did not need conditioning. Animals 2, 3, and 4 were immediately comfortable in the chamber after recovering from sedation for electrode connections. Animals 5 and 6 had two conditioning sessions exceeding 30 minutes in the chamber either after (Animal 5) or before surgery (Animal 6). During testing in the chamber, touch, wet food, toys, and even animal videos were used to help maintain a calm animal, as needed.

An important part of minimizing stress during awake sessions, and thus allowing more testing time, was to minimize the strain exerted from the cables to the backpack. We developed a more refined setup which used lightweight stimulation cables and neural signal headstage cables and bundled them with the two lightweight bladder catheters with twist ties. We left enough length for the animal to move around in the testing area and budgeted extra length for animal twisting and turning. We also transitioned from a large encircled testing area used in our prior behavioral study [2] to a smaller awake testing cage with a meshed floor to monitor urine flow and allow for easier handling (Figure 1) [3].

From Animals 1 to 3, we found a few sources that introduced noise to the bladder pressure and DRG microelectrode signals and developed strategies to mitigate their effects. We first learned that maintaining a normal body temperature is crucial under alfaxalone. A high level of noise on the bladder pressure line can be induced by even the lowest level of animal shivering due to low body temperature as an animal recovers from alfaxalone sedation. Other types of moving (e.g., walking, eating, turning) are admittedly another considerable source of signal artifacts, which we addressed to a limited degree using a cross-channel invalidation method as a DRG microelectrode signal processing step. It is also possible that electromagnetic interference compromised the signal quality, but we only observed it a few times in the experiments and did not consider it a major concern. We wrapped the backpack and headstages with aluminum foil for a few early sessions, which seemed to help mitigate this occurrence although we did not continue the practice in later Animals.

To improve experimental efficiency, we increased the bladder infusion rate from 2 ml/min (all of Animal 1, most of Animal 2, and some of Animal 3) to 5 ml/min (some of Animal 2 and most of Animal 3). Although this higher rate is well above physiological urine generation [4], we did not observe an effect on bladder capacity. In Animal 3, the capacity for 2 ml/min (37.1 ± 8.7 ml, n = 29 trials) was essentially the same as for 5 ml/min (37.3 ± 8.0 ml, n = 56 trials), across a mix of awake and sedated trials with and without stimulation. In some sessions, using a 5 ml/min infusion rate led to infusion pump artifacts on the bladder pressure. We did not consider them a concern, as the artifact amplitudes were very low (~1–2 cmH2O). Continuing, we used 5 ml/min in Animal 4, mostly 5 ml/min in Animal 5 (with some 2 ml/min trials), and 2 ml/min in Experiment 6 due to a small bladder volume.

#### Part 2 – Tables and Figures

**Table S1.** Stimulation electrode functionality across Animals.

| Experiment (Animal) | Sedated sessions | Awake sessions (joint w/ sedated) | Sacral threshold range (mA) | Sacral electrode duration (days) | Pudendal threshold range (mA) | Pudendal electrode duration (days) |
|---|---|---|---|---|---|---|
| 1 | 7 | 3 (3) | 0.20-0.32 | 13 | 0.09-0.64 | 33# |
| 2 | 6 | 8 (5) | 0.09-0.21 | 63# | NA | 0 |
| 3 | 9 | 13 (9) | 0.23-1.50 | 21 | 0.13-0.66 | 56# |
| 4 | 5 | 7 (5) | 0.3 | 1 | 0.17-0.31 | 92# |
| 5 | 5 | 7 (2) | 0.15-4.10 | 63# | 0.19-0.55 | 63# |
| 6 | 2 | 7 (2) | 0.07-0.14 | 70# | 0.10-0.14 | 70# |
| 7 | 0 | 0 | 0.02-0.03 | 5# | 0.09-0.13 | 5# |

### - indicates electrode still functional at experiment end-date duration

**Table S2.** Count of identified DRG units across Animals 2-7 during behavioral testing. No behavioral testing was performed in the Animal 1 awake sessions. No units were detected in Animal 5.

| Animal | Bladder units | Bladder unit correlation to pressure | Putative flow units | Brushing or movement units | Other units | Total units |
|---|---|---|---|---|---|---|
| 2 | 28 | 0.31 ± 0.10 (0.20-0.61) | 1 | 1 | 69 | 99 |
| 3 | 23 | 0.41 ± 0.14 (0.20-0.74) | 4 | 3 | 13 | 43 |
| 4 | 27 | 0.49 ± 0.21 (0.21-0.92) | 1 | 5 | 31 | 64 |
| 6 | 0 | -- | 0 | 1 | 7 | 8 |
| *Mean* | *19.5* | 0.40 ± 0.19 | *1.5* | *2.5* | *30* | *53.5* |

**Table S3.** Summary of litterbox visits across animals per 24-hour periods. Animal 2 did not leave the recovery cage, and Animals 4 and 5 stayed in the cage longer during recovery. Values are given as mean ± standard deviation (count). Statistical differences across groups are indicated in Figure S6A.

| Animal | Pre-Surgery | Post-Surgery | Pre-Session | Post-Session | Animal Total |
|---|---|---|---|---|---|
| 1 | 4.4 ± 1.3 (7) | 63.8 ± 38.8 (4) | 40.3 ± 21.7 (8) | 35.6 ± 14.7 (6) | 36.1 ± 13.5 |
| 3 | 3.7 ± 0.8 (7) | 9 (1) | 15.4 ± 6.3 (7) | 16.4 ± 5.4 (7) | 11.1 ± 2.8 |
| 4 | 2.0 ± 0.8 (7) | *in cage* | 7.1 ± 3.7 (6) | 7.0 ± 3.2 (5) | 7.0 ± 3.3 |
| 5 | 2.1 ± 0.7 (7) | *in cage* | 7.6 ± 8.3 (5) | 8.4 ± 8.3 (5) | 6.1 ± 3.6 |
| 6 | 3.7 ± 1.3 (7) | 5.4 ± 2.4 (5) | 43.9 ± 24.3 (16) | 41.5 ± 15.3 (16) | 23.6 ± 9.5 |
| Per-Animal Summary | 3.2 ± 1.1 (5) | --- | 22.2 ± 18.7 (5) | 21.0 ± 16.8 (5) | |

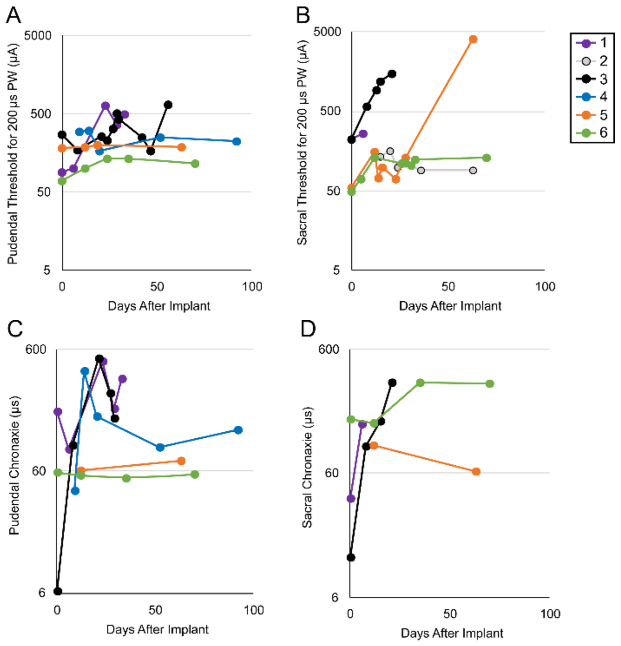


**Figure S1.** Stimulation electrode functionality across Animals. At top, stimulation threshold levels for anal sphincter visual twitch responses are given for A. pudendal nerve and B. sacral nerve stimulation across experiment durations. Stimulation pulse widths at both 200 and 210 µs are combined. The sacral nerve electrodes had more threshold variability than the pudendal nerve electrodes. This was likely due to structural differences between the electrodes. At bottom, chronaxie values from strength-duration curves for C. pudendal nerve and D. sacral nerve stimulation are given across experiment durations. Strength-duration responses were not obtained in all Animals and sessions. We believe variations in chronaxie and thresholds over time were attrituable to changes in electrode stability (e.g., scar tissue formation, electrode movement) as compared to changes in nerve excitability (e.g., nerve damage, neuroplasticity). These variations did not have an impact on bladder outcome measures.

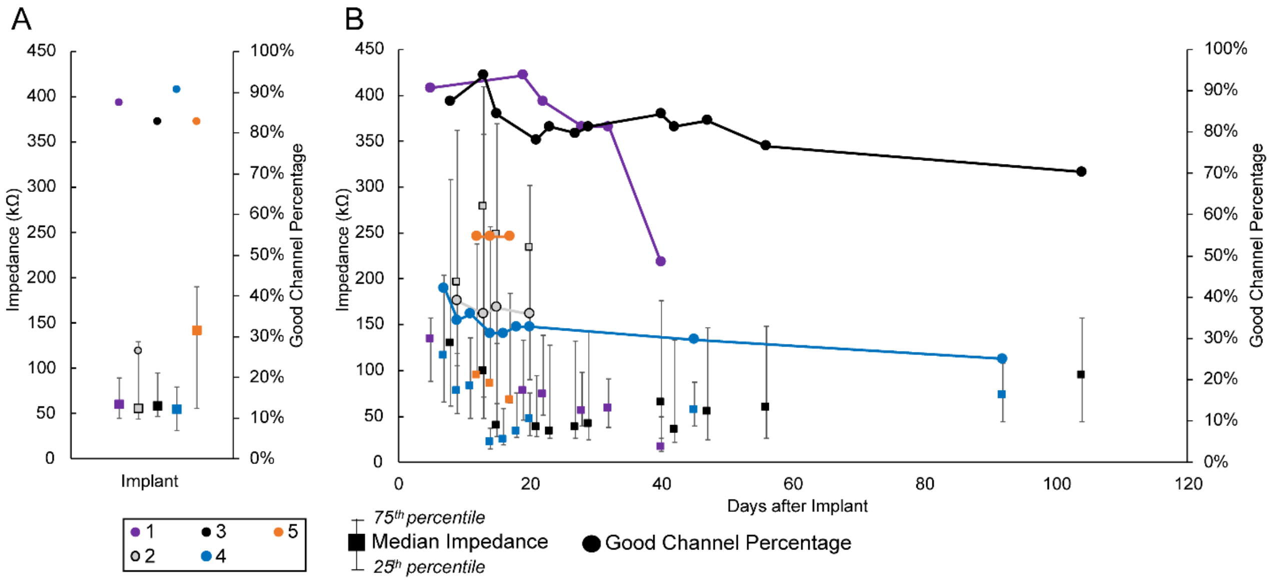


**Figure S2.** Summary of DRG electrode functionality. A. Median microelectrode impedances at 1 kHz during implant surgery for "good channels" (impedance < 1 MΩ), on the left axis (squares). Percentage of channels with good impedance across Animals on right axis (circles). B. Median microelectrode impedances (squares; left y-axis) and good channel percentages (connected circles; right y-axis) after the implant surgeries.

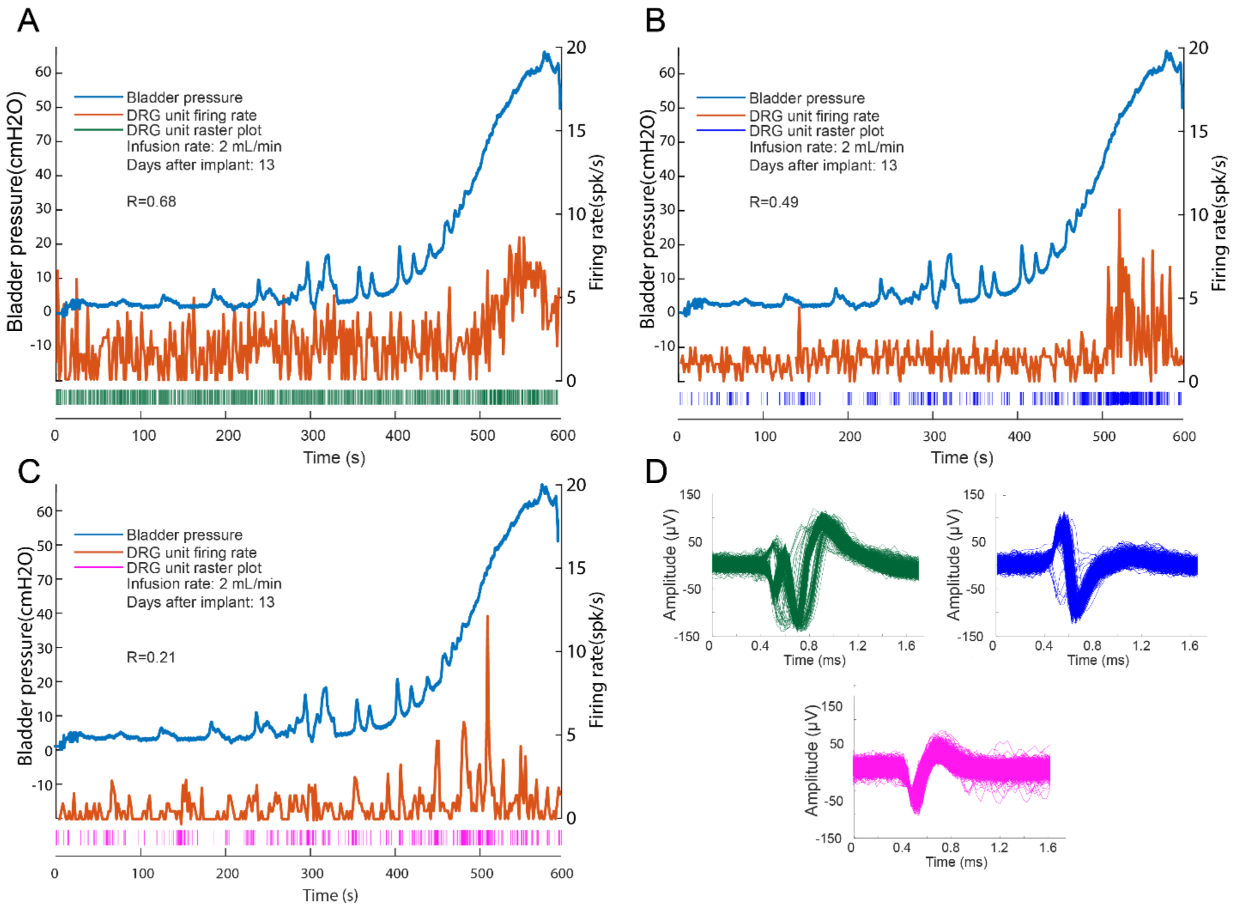


**Figure S3.** Example DRG bladder units during an awake no-stimulation bladder fill in Animal 3, 13 days after implant. A-C. The firing rate and raster for three DRG bladder units are shown with the bladder pressure. The firing rate was calculated using a 1-second window without overlap. D. Waveforms for each bladder unit (waveform color is matched to the corresponding raster plot). The units correspond to unit 60.1 (green), unit 64.2 (blue), and unit 7.1 (pink) in Figure S5A.

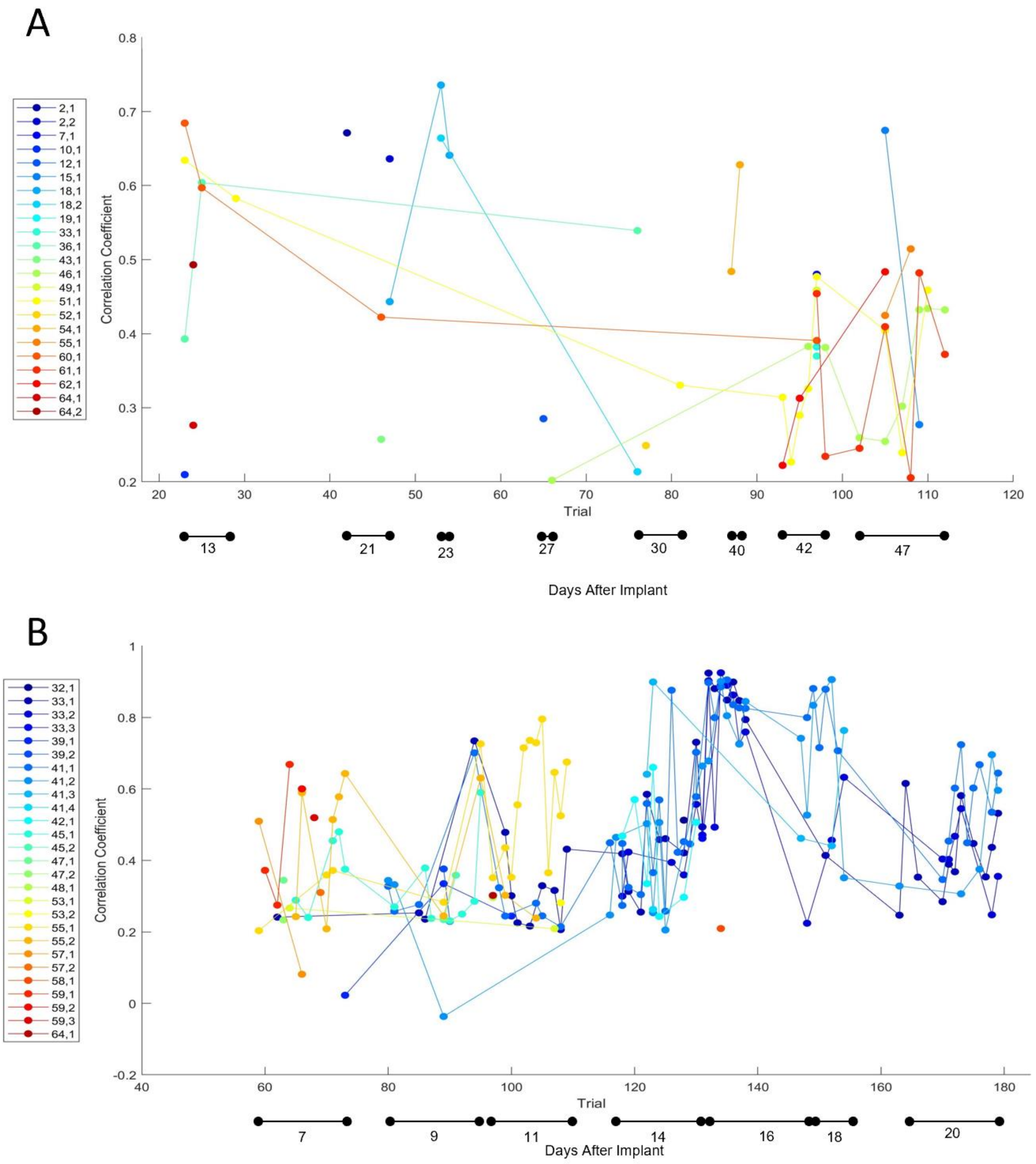


**Figure S4.** Sorted single units with firing rate correlation coefficient > 0.2 to bladder pressure for at least one test session in A. Animal 3 and B. Animal 4 across testing periods. Trial numbering on the y-axes is inclusive of all trials with data collected, including non-bladder trials such as impedance checks and stimulation electrode testing; bladder units were only identified during bladder test trials. In Animal 3, unit 51.1 was the longest tracked unit, at 34 days from day 13 to day 47. In Animal 4, unit 33.1 was tracked the longest, 13 days from day 7 to day 20. Of the 27 units in Animal 4, 11 appear in more than one session, with 33.2 (4), 41.2 (5), 41.1 (6) and 33.1 (7) having the most active sessions.

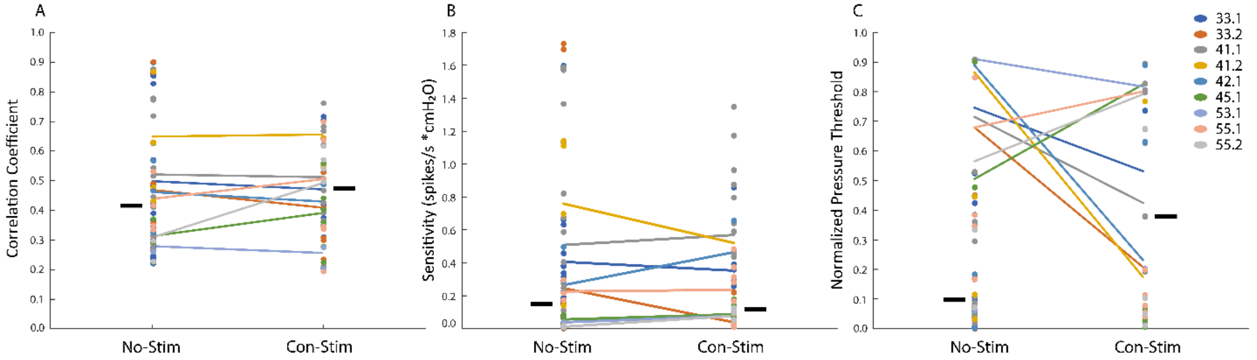


**Figure S5.** Bladder unit relationships to bladder pressure did not significantly vary between no-stimulation and continuous stimulation trials in Experiment 4. A. The correlation coefficient of DRG bladder unit firing rates to bladder pressure (Welch's t-test, p-value = 0.92). B. The linear regression slope (sensitivity) for DRG bladder unit firing rates to bladder pressure (Welch's t-test, p-value = 0.70. C. The bladder pressure threshold for DRG bladder unit firing normalized to the bladder pressure when voiding occurs (Welch's t-test, p-value = 0.09). Non-grey icons designate units identified across multiple trials, with units present in sequential no-stimulation and continuous stimulation trials connected with a line. Horizontal black bars indicate overall group medians.

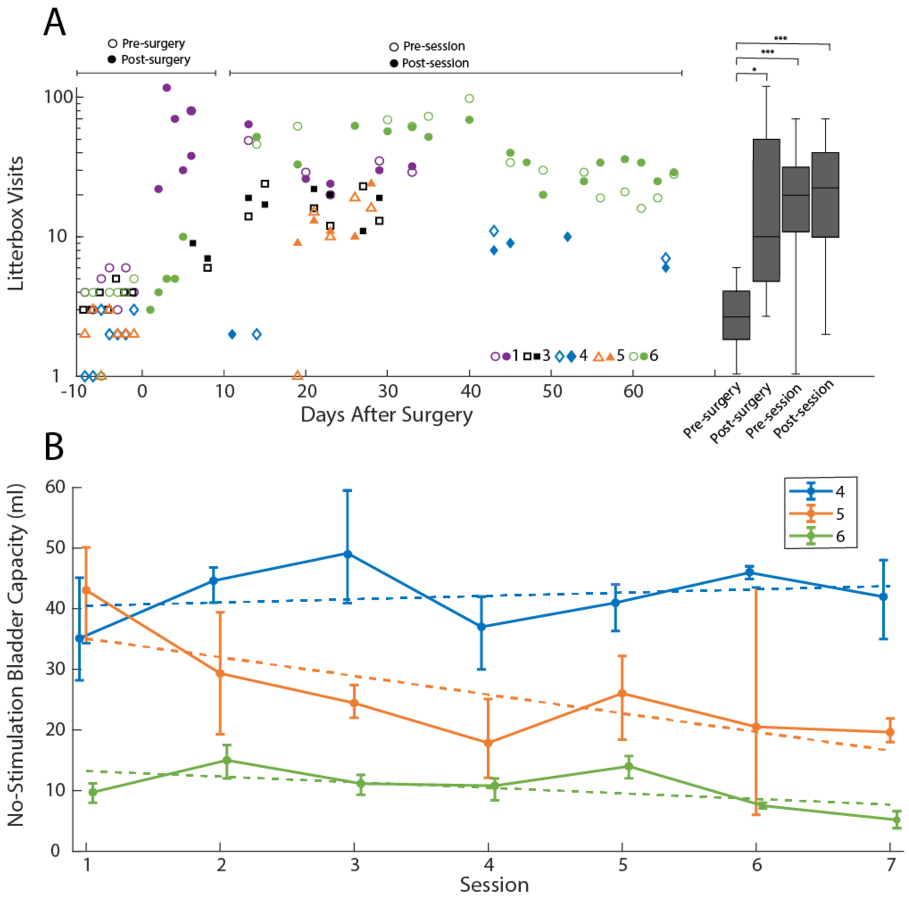


**Figure S6.** A. Analysis of feline litterbox use during study. The scatterplot contains 24-hour interval counts of litterbox usage before surgery (open icons), within ~one week after surgery (closed icons), and before (open icons) and after (closed icons) test sessions. The boxplots summarize the litterbox visits per period across animals and are organized as in Fig. 2. Animals 3 and 4 spent most of the time within a recovery cage in the post-surgery period. Animal 2 spent his entire implant time in the cage and is omitted here. Some pre- and post-session periods for Animal 4 had no litterbox visits. (* $p < 0.05$. *** $p < 0.001$) B. Average no-stimulation bladder capacities for animals with repeat test sessions. Dashed lines are linear regression fits. (slope p-values: Animal 4 = 0.73, Animal 5 = 0.001, Animal 6 = 0.002).
